# Striatal M4 Receptors Mediate M4 PAM Antipsychotic-like Efficacy Independently of Normalizing Striatal Dopamine Release

**DOI:** 10.64898/2026.10.26.761976

**Authors:** Daniel G Joyner, Curran M Johnston, Yumin Choi, Michael N May, Rebecca D Carden, Daniel J. Foster

## Abstract

Muscarinic acetylcholine receptor (mAChR) drugs have emerged as a promising new class of antipsychotics, highlighted by the FDA approval of Cobenfy, the first treatment for schizophrenia that does not directly target dopamine receptors. The M4 receptor subtype is thought to be a key mediator of antipsychotic-like efficacy. However, the *in vivo* circuit mechanisms underlying the antipsychotic-like efficacy observed with M4-selective positive allosteric modulators (PAMs) remain incompletely understood. Here, we used fiber photometry in awake, behaving mice to assess how the M4 PAM VU0467154 (VU’154) affects striatal dopamine (DA) transmission under basal and amphetamine-stimulated conditions. We also generated and validated dorsal striatum-specific M4 knockout mice (DS-M4-KO) to test the role of striatal M4 receptors in regulating DA release and antipsychotic-like effects. VU’154 dose-dependently reduced the amplitude of striatal DA release events under basal conditions without altering event frequency. However, VU’154 had no significant effects on tonic DA levels under basal conditions, as assessed using the D1 PAM DETQ. Despite robustly attenuating AHL, VU’154 produced little to no effect on amphetamine-induced DA levels, indicating that antipsychotic-like efficacy was dissociated from changes in striatal DA levels. Notably, VU’154 efficacy in AHL was absent in DS-M4-KO mice, demonstrating that M4 receptors in the dorsal striatum are required for antipsychotic-like effects. Combined with previous studies implicating a key role for M4 receptors on D1-expressing neurons, these findings suggest that M4 PAMs produce antipsychotic-like efficacy primarily through postsynaptic mechanisms in the dorsal striatum, rather than through broad normalization of amphetamine-induced DA release.

**Graphical Abstract:** 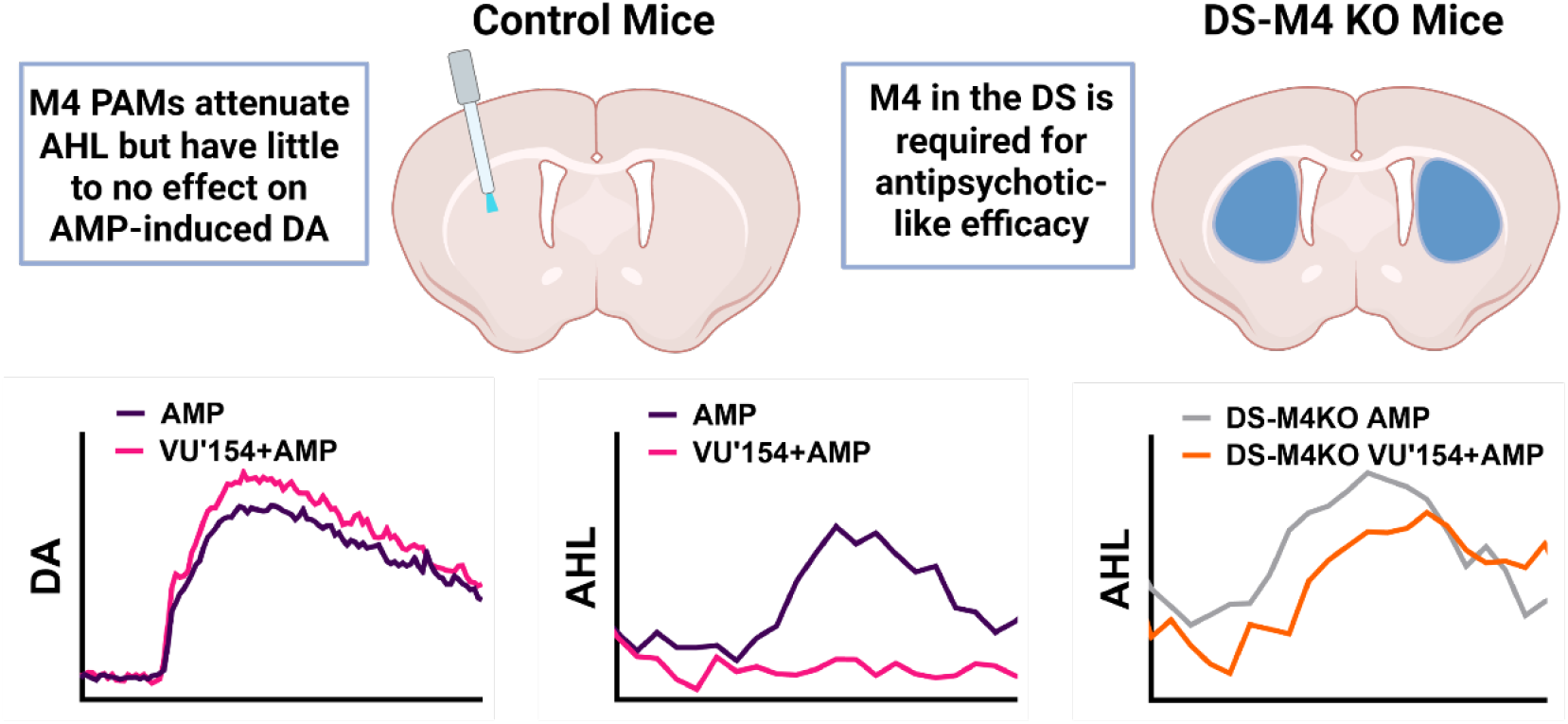

## Introduction

Schizophrenia is a complex disorder associated with numerous physiological changes in the brain [1–3], with dysregulated dopamine (DA) signaling among the most consistently implicated mechanisms [4–6]. While the dopamine hypothesis has been refined over the years, our current understanding posits that there is hypodopaminergic transmission to the prefrontal cortex, a potential contributor to cognitive deficits [5, 7], while transmission in the dorsal striatum (DS) is hyperdopaminergic [8], which seems to play a key role in positive symptoms of the disorder. It is the latter feature through which D2 antagonists are thought to confer a majority of their antipsychotic efficacy, while the former is likely why these medications do little for, or even exacerbate, cognitive deficits [9–12]. The lack of efficacy in treating negative symptoms and cognitive deficits, along with the numerous side effects observed with current treatments, highlights an urgent need for novel therapeutic strategies [13–15].

Muscarinic acetylcholine receptors (mAChRs) have garnered increasing attention for their antipsychotic potential, with Cobenfy recently becoming the first mAChR-targeting drug approved by the FDA for schizophrenia [16, 17]. Preclinical studies using receptor subtype-selective drugs and knockout mice suggest that the muscarinic M1 and M4 receptors are the primary mediators of pro-cognitive and antipsychotic-like effects observed with mAChR-based interventions [18–25]. M4-specific compounds display antipsychotic-like efficacy in several preclinical assays, including both attenuation of amphetamine-induced hyperlocomotion (AHL) and disruption of pre-pulse inhibition [19, 26, 27]. Given the link between hyperdopaminergic striatal signaling and positive symptoms of schizophrenia, and the fact that M4 receptors are densely expressed in the striatum [28–30], an M4 receptor-mediated reduction in DA has been hypothesized to be a potential mechanism mediating muscarinic antipsychotic efficacy. In support of this, non-selective muscarinic agonists have been shown to reduce electrically evoked DA release in brain slices, with M4-specific positive allosteric modulators (PAMs) enhancing this effect [26, 31, 32]. In addition to attenuating AHL, M4 PAM pretreatment reduces amphetamine-induced increases in brain activity in the striatum and other brain regions [19]. However, our understanding of how muscarinic drugs regulate neurotransmission *in vivo* remains limited, including precisely which M4 receptor population confers antipsychotic-like efficacy.

Here, we address this gap using fiber photometry approaches in awake-behaving mice and determine the ability of M4 PAM VU0467154 (VU’154) to alter striatal DA transmission under both basal and amphetamine-induced conditions. We then use region-specific M4 knockout mice to rigorously assess the importance of M4 receptors expressed in the DS in regulating striatal signaling and mediating antipsychotic-like efficacy. Collectively, these studies provide key insights into the mechanisms whereby mAChR-targeting drugs regulate schizophrenia-associated circuitry and mediate antipsychotic-like efficacy.

## Material and Methods

### Animals

Experiments utilized both male and female mice that were group-housed and provided with *ad libitum* food and water on a 12-hour light/dark cycle (lights on at 7:00 am). Wildtype (Jackson # 000664 WT), DAT-Cre (Jackson strain # 006660), and M4^fl/fl^ [20, 26] mice were used as outlined below. All experiments were approved by the University of South Carolina Institutional Animal Care and Use Committee. Stereotaxic surgery was performed in 9–12-week-old mice and experiments outlined below were performed 3-5 weeks following surgery to allow time for virus expression.

### Drugs

VU0467154 (VU’154) was generously provided by the Vanderbilt Center for Neuroscience Drug Discovery and was dissolved in vehicle (10% Tween 80). D-amphetamine hemisulfate (Sigma-Aldrich) was dissolved in sterile saline. DETQ [2-(2,6-dichlorophenyl)−1-((1S,3R)−3-(hydroxymethyl)−5-(2-hydroxypropan-2-yl)−1-methyl-3,4-dihydroisoquinolin-2(1H)-yl)ethan-1-one] (Hello Bio, SKU HB15406) was dissolved in 20% βCD (20% w/v hydroxypropyl beta cyclodextrin (Cyclodextrin Technologies)). All drugs were administered at 10 mL/kg.

### Stereotaxic Surgery

#### Spontaneous and amphetamine-hyperlocomotion experiments

Mice received 0.3 µL of dLight1.3b (pAAV5-CAG-dLight1.3b, Addgene 125560-AAV5) unilaterally injected into the dorsal striatum (DS; coordinates, AP: +1.2, ML: ±1.8, DV: −3.0) at 0.05 µL/min. Then, a 1.25 mm fiber optic ferrule cannula (0.48NA, 3.0mm long, 400μm core; Doric) was implanted and secured to the skull using dental cement (for additional details see Supplementary Methods).

#### Viral-induced M4-KO

M4^f/f^ mice received bilateral injections of 0.15-0.25 µL of Cre recombinase (pENN.AAV5.CamKII 0.4.Cre.SV40, Addgene 105558-AAV5) or 0.15-0.25 µL of mCherry (pAAV5-CamKIIa-mCherry, Addgene 114469-AAV5) into the DS at 0.05 µL/min, followed by unilaterally injection of dLight1.3b and cannulation as described above.

### Fiber Photometry

Photometry experiments utilized an RZ10X Lux-I/O Processor (Tucker-Davis Technologies) using integrated LEDs (465 nm experimental and 405 nm control). Light intensities used were 20–35 µW for 465 nm and 5–12.5 µW for 405 nm. Photometry data were processed in MATLAB (MathWorks) using custom code. Briefly, data streams were low-pass filtered and a linear least-squares fit was generated by fitting the 405 signal to the 465 signal, with the resulting fitted signal being used to calculate ΔF/F. Spontaneous event transients were detected as transients at or exceeding 2.91 times the median absolute deviation (MAD), as adapted from previously published methodology [33]. dLight1.3 signals in the presence of dopamine receptor PAM (DETQ) were based on previously published approaches using the 10 min period prior to drug administration as the baseline for normalization. dLight1.3 signals in the presence of amphetamine were based on previously published approaches [34] using the 30 min period prior to drug administration served as the baseline for normalization. All custom code used is available on GitHub (https://doi.org/10.5281/zenodo.23290497, see Supplemental Methods for additional details).

#### Assessment of Basal Striatal Dopamine Release and Locomotion

These experiments utilized a within-subjects design. Mice were attached to the fiber-photometry patch cable, placed in the arena, and recorded prior to and following administration of VU’154 (0-3 mg/kg; i.p.). In experiments assessing tonic changes in DA signaling, DETQ (10 mg/kg; i.p.) was administered at the same time as vehicle (10% Tween 80; i.p.) or VU’154 (3 mg/kg; i.p.). All trials were run with 48 hours between doses to allow for drug washout. Each mouse received a semi-randomized dose/condition each session and was exposed to all doses/conditions once over the course of the experiment.

#### Amphetamine Hyperlocomotion Experiments

All mice were amphetamine-naïve to avoid confounding effects of prior stimulant exposure. Fiber-photometry and video recordings were acquired simultaneously. Mice were allowed to freely explore the open-field arena for 90 min before receiving an injection of either VU’154 (3 mg/kg, i.p.) or vehicle (10% Tween 80, i.p.). Mice were then returned to the arena for 30 min before receiving D-amphetamine (3 mg/kg, s.c.) and were then recorded for an additional 2 hours.

### Tissue Processing

#### Immunohistochemistry

Mice were perfused with 4% paraformaldehyde (PFA), sectioned at 40-50 µm on a freezing microtome. dLight1.3 expression was visualized by immunostaining with chicken anti-GFP primary antibody (1:4000; ab13970, Abcam) followed by goat anti-chicken Alexa Fluor 594 secondary antibody (1:500; ab150176, Abcam; See Supplemental Methods for details).

#### RNAscope

Brains were removed, flash-frozen, sectioned coronally (20 µm) on a cryostat, and assessed using The Advanced Multiplexing RNAscope V2 Assay (Advanced Cell Diagnostics). Briefly, probes (Mm-Chrm1-O1, Mm-Chrm4-C2, Mm-Chat-C3) were conjugated to TSA Vivid 570 (yellow, 1:4000), TSA Vivid 650 (red, 1:5000), and TSA Vivid 520 (green, 1:3000), respectively, and imaged on a Leica Stellaris 5 Confocal Microscope. To quantify M1-and M4-positive cells, the channels were analyzed using QuPath, Fiji, and custom script (https://doi.org/10.5281/zenodo.23301431; see Supplemental Methods for details).

### Locomotion Analysis

DeepLabCut was used to assess locomotion as previously reported [35]. Distance traveled was calculated from the centroid position and total distance traveled and time in the center of the open field were calculated (see Supplemental Methods for details).

### Statistical Analysis

Dose-response data were expressed as % inhibition relative to baseline and analyzed via a repeated measures one-way ANOVA to determine if there was a significant effect of treatment (α = 0.05). Sex differences were assessed via two-way ANOVA to determine main effects of sex and treatment and to assess sex x treatment interactions. For DETQ experiments, amplitude % change from baseline and area under the curve (AUC) were analyzed using paired t-tests (α = 0.05). AUC values for dLight signals, locomotion activity during AHL, as well as baseline locomotor activity and time spent in the center of the open-field were analyzed using unpaired t-tests (α = 0.05). Data were analyzed using Prism 11 (GraphPad Software).

## Results

### The M4 PAM VU’154 dose-dependently reduces striatal DA release event amplitudes

We first set out to assess the ability of the M4 PAM VU’154 to dose-dependently regulate striatal dopamine release *in vivo*. Animals were unilaterally transduced with the dLight1.3 sensor and fiber-optic cannulas were placed in the DS (**Fig. 1A-C**). Detected DA release events were validated by control experiments in which DA transients were blocked by a DA receptor antagonist (**Fig. S1C**) or induced by optogenetic activation of DA neuron terminals (**Fig. S1D,E**). Following a 10 min baseline period, mice were dosed with VU’154 (0, 0.3, 1, and 3 mg/kg, i.p.) and DA event amplitudes and frequencies were assessed. Treatment with vehicle (10% Tween-80; i.p.) had little to no effect on DA event frequency or amplitudes (**Fig. 1D, S1B**).

**Figure 1:**
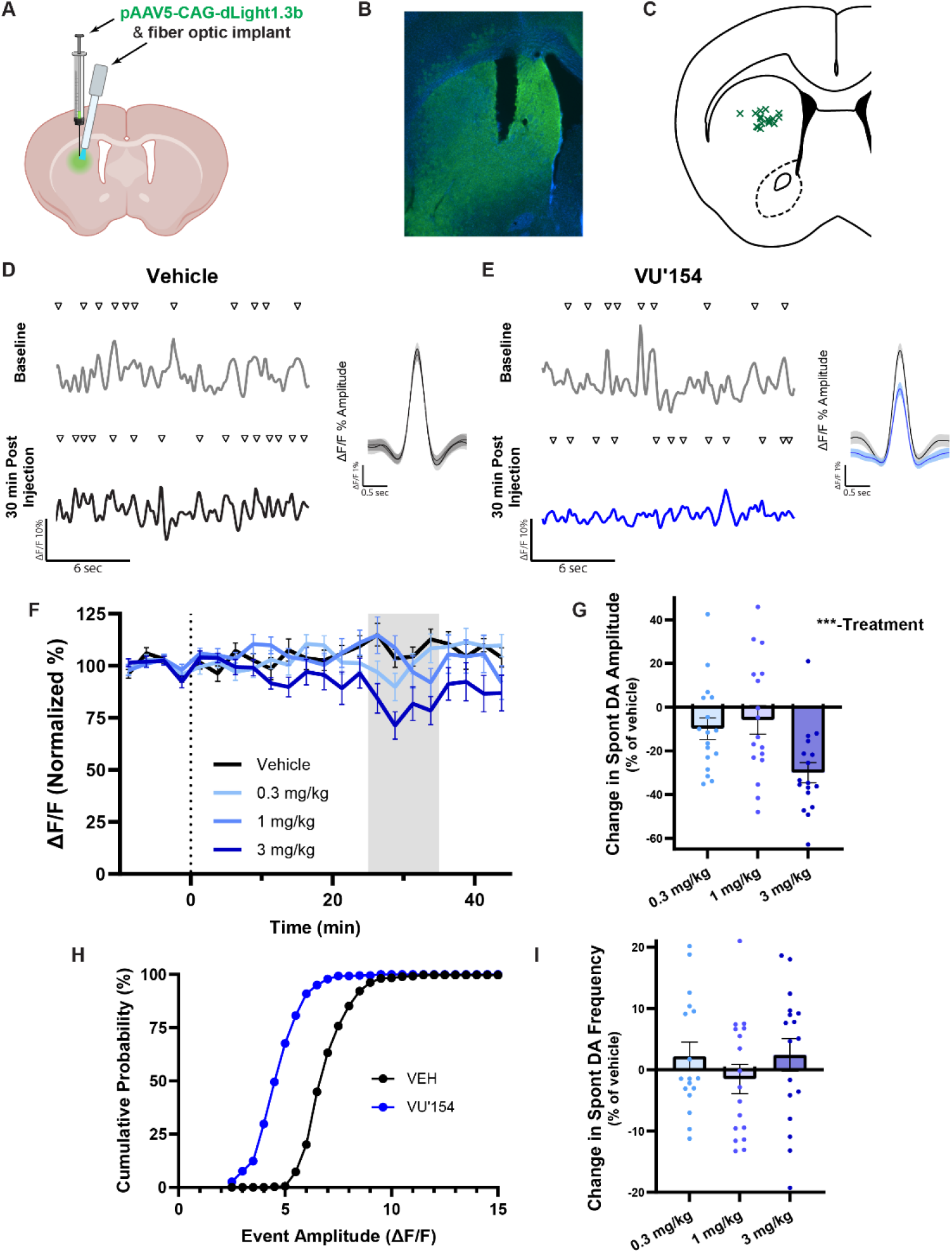
M4 PAM VU’154 dose-dependently reduces striatal DA release event amplitudes. (**A**) Diagram depicting injection of dLight virus and fiber optic cannula implantation. (**B**) Immunofluorescent image showing cannula placement and dLight expression in dorsal striatum. (**C**) Verified cannula placements of WT mice (n = 17). (**D**) Representative snippets with detected spontaneous DA release events and mean composite trace (SEM 95% CI) at baseline and post-injection, in vehicle and (**E**) 3 mg/kg of VU’154. (**F**) Time course of dose response, normalized to baseline, dotted line = injection, gray area = “peak period”, 25-35 min post-injection, used for analysis. (**G**) Change in event amplitudes relative to vehicle showing main effect of treatment, presented as mean ± SEM, (one-way ANOVA, p < 0.0005). (**H**) Cumulative probability of events by amplitude during peak period. **(I**) Change in event frequency (one-way ANOVA, p > 0.05).

However, administration of 3mg/kg VU’154 was able to cause a robust inhibition of DA event amplitudes (**Fig. 1E**), with a time course exhibiting peak effects ∼30 min post-dosing (**Fig. 1F**). Repeated measures one-way ANOVA showed a significant main effect of treatment (n = 18, p = 0.0005) to reduce DA release amplitudes (**Fig. 1G**). To assess if the VU’154 effects were specific to certain subsets of detected events, we examined cumulative probability distributions of event amplitudes. The approximately parallel shift across the distribution (**Fig. 1H**) indicates that VU’154 affected both small-and large-amplitude events. While VU’154 reduced event amplitudes, there was no main effect of treatment on event frequency (p = 0.185, **Fig. 1I**). No main effect of sex was detected (2-way ANOVA, treatment: p = 0.002, sex: p = 0.69, sex by treatment: p = 0.313) (**Fig. S1A**) and accordingly, all the data assessing DA event amplitudes shown moving forward are presented with sexes combined.

### Assessment of VU’154 effects on tonic DA levels

Due to its pharmacological profile, DETQ can enhance DA binding to dLight, and thus improve the signal-to-noise ratio of dLight-mediated photometry recordings [36]. Administration of DETQ (10 mg/kg, i.p.) caused a robust enhancement of dLight signal as assessed by an upward shift in both Z-score (**Fig. 2A**) and ΔF/F (**Fig. 2C**). Consistent with the ability of DETQ to bind dLight and not to mouse D1 receptors, this DETQ-mediated increase in dLight signal was not associated with an increase in locomotion (**Fig. S2**). When 3 mg/kg VU’154 was co-administered with DETQ, the magnitude of DETQ effects on z-score were not significantly different from those obtained with vehicle and DETQ (n = 12, paired t test, p = 0.242, **Fig. 2A,B**), suggesting that VU’154 has little to no effect on tonic DA levels. When DETQ was applied, it not only changed the overall dLight fluorescence, but also increased the amplitude of detected DA events (165.7 ± 13.2% of baseline; **Fig. 2C,E**), and VU’154 co-administration significantly reduced the magnitude of this increase (130.2 ± 5.7 % of baseline, paired t-test, p = 0.018, **Fig. 2G**). Similar to results obtained in the absence of DETQ, VU’154 co-administered with DETQ caused an evenly distributed reduction in event amplitudes (**Fig. 2H**), without altering event frequency (**Fig. 2I**). Collectively, these results suggest that M4 activation can inhibit DA transient amplitudes while having little to no effect on tonic DA levels.

**Figure 2:**
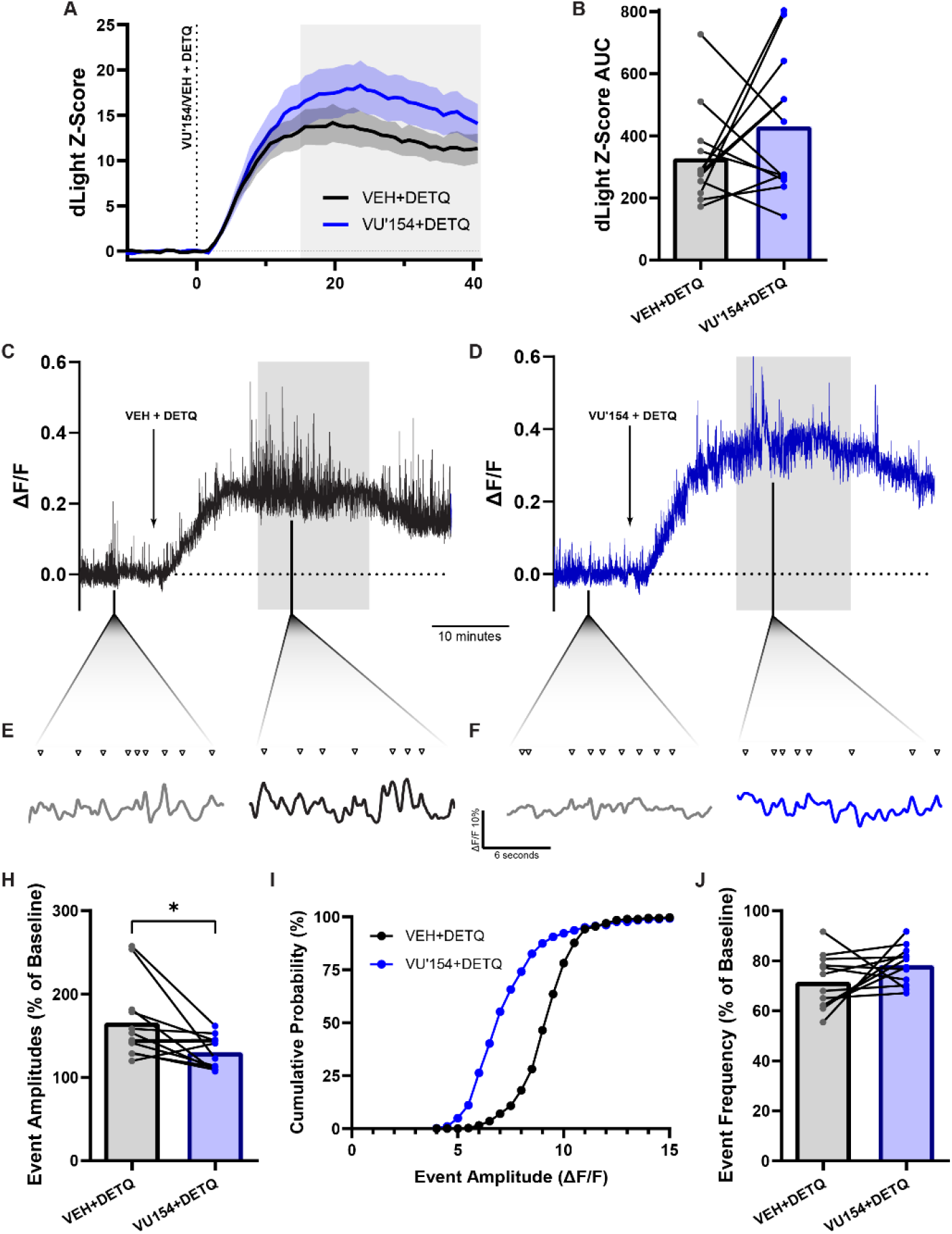
Assessment of VU’154 effects on tonic DA levels. (**A**) Time course of change in measured dLight fluorescence intensity (Z-score relative to 10 min pre-injection). (**B**) Averaged data depicting dLight Z-score AUC (paired t-test, p > 0.05). (**C**) Representative trial trace following coadministration of DETQ (10 mg/kg i.p.) with vehicle and (**D**) 3 mg/kg of VU’154. (**E**) Snippets of trace labeled with detected events and baseline and peak period after DETQ with vehicle and (**F**) 3 mg/kg of VU’154. (**G**) Averaged data depicting mean event amplitudes at peak period relative to baseline (within subjects) (paired t-test, p < 0.05). (**FH**) Cumulative probability of events by amplitude at peak period after VEH or VU1’54. (**I**) Averaged data depicting mean event frequency at peak period (paired t-test, p > 0.05).

### VU’154 induces antipsychotic-like effects while potentiating amphetamine-induced striatal DA release

M4 PAMs have previously been reported to induce antipsychotic-like efficacy in the amphetamine-induced hyperlocomotion assay (AHL) [19, 37]. Here, to elucidate the mechanisms behind this efficacy, we combined the AHL assay with fiber photometry to characterize VU’154 effects on amphetamine-induced DA changes in the DS. Mice were pretreated with vehicle or VU’154 (3 mg/kg; i.p.) 30 min prior to administration of amphetamine (3 mg/kg; s.c.; **Fig. 3A**) and DA levels were tracked via fiber photometry recordings while locomotion was tracked using DeepLab Cut (**Fig. 3B**). As previously observed, pretreatment with M4 PAM caused a significant reduction in AHL compared to mice pretreated with vehicle (n = 6-7, unpaired t-test, p = 0.007; **Fig. 3C,D**). VU’154-mediated reductions in locomotion were not accompanied by reductions in amphetamine-induced DA levels, as VU’154 treatment significantly increased amphetamine-induced DA levels compared to mice pretreated with vehicle (n = 6-7, unpaired t-test, p = 0.024; **Fig. 3E,F**). Further analysis did not reveal any sex-specific effects on either AHL or dLight (**Fig. S3**, 2-way ANOVA, dLight: Treatment: p = 0.028, Sex: p = 0.113. AHL: Treatment: p = 0.012, Sex: p = 0.535). Taken together, these studies suggest that while VU’154 can robustly reduce AHL, these effects are not through normalization of striatal DA levels.

**Figure 3:**
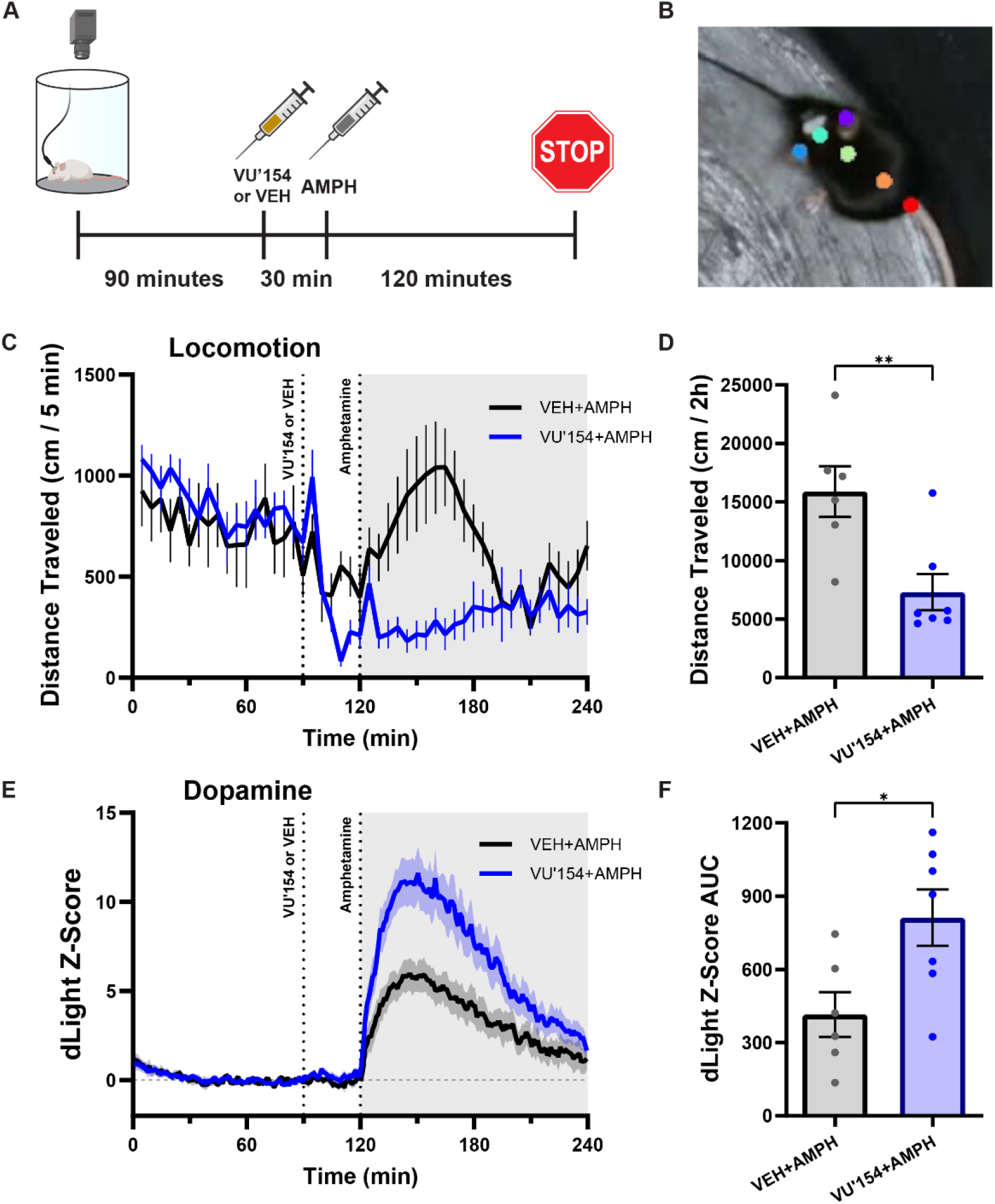
VU’154 induces antipsychotic-like effects while potentiating amphetamine-induced striatal DA release. (**A**) Experimental timeline. (**B**) Image from DeepLabCut-labeled video. Orange ‘Centroid’ was used to analyze locomotion. (**C**) Time course plotting distance traveled per 5-min bin. Gray area used for AUC. (**D**) Averaged data depicting a significant attenuation of hyperlocomotion in WT mice pretreated with 3 mg/kg VU’154 (unpaired t-test, p < 0.005). (**E**) Time course of change in dLight fluorescence in WT mice. (**F**) Averaged data depicting a significant difference in dLight AUC post-AMPH (unpaired t-test, p < 0.05).

### Selective deletion of M4 receptors from the dorsal striatum attenuates VU’154 antipsychotic-like efficacy

Previous studies indicate that M4 receptors expressed on D1-expressing spiny projection neurons (D1-SPNs) play a key role in mediating antipsychotic-like efficacy [26, 37–39]. To further test this hypothesis, we deleted M4 from the DS (DS-M4-KO) using viral approaches. Transgenic M4^fl/fl^ mice were bilaterally injected in the DS with an AAV encoding either Cre-recombinase or mCherry (**Fig. 4A**). In the nucleus accumbens (NAc), we observed robust expression of M1 and M4 receptor RNA, with M1 RNA expression observed in 94.1% of cells and 45.8% of those M1-positive cells also expressing M4 RNA (**Fig. 4D,E**). However, M4 RNA was almost completely ablated in the DS of mice receiving AAV encoding Cre-recombinase (1.1% of cells expressing M4 RNA) with little to no change in M1 receptor RNA expression (**Fig. 4B-E**). M4 RNA expression was lost broadly across the DS, spanning both the dorsomedial and dorsolateral areas, but did not extend into the NAc (**Fig. 4B**). The reduction in striatal M4 RNA expression was not restricted to SPNs, as M4 receptor RNA was also observed to be deleted from cholinergic interneurons of the DS (**Fig. S4A,B**). Similar to results in global M4 knockout mice or D1-M4 KO mice [20, 40, 41], region-specific deletion was associated with an overall hyperlocomotive phenotype when compared to mCherry controls (**Fig. S4C**). Together these results demonstrate a selective and region-specific deletion of M4 from the DS.

**Figure 4:**
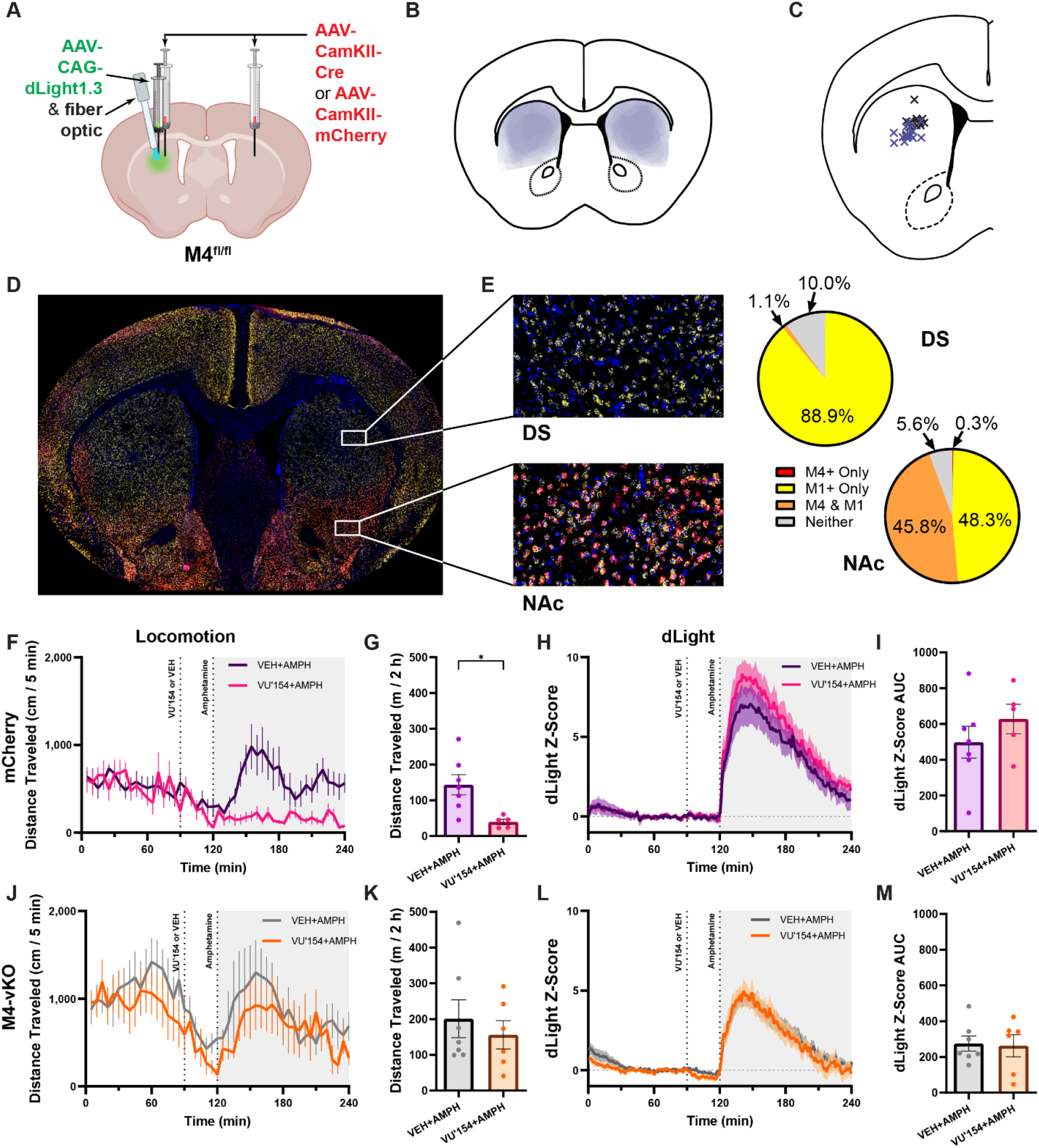
Selective deletion of M4 receptors from the dorsal striatum attenuates VU’154 antipsychotic-like efficacy. (**A**) Diagram depicting viral injections, bilateral Cre or mCherry, then unilateral dLight and fiber optic cannula implantation. (**B**) Diagram depicting overlays of M4 deletion area, fluorescent images fitted to brain atlas. (**C**) Verified cannula placements of DS-M4-KO mice. (**D**) Representative RNAscope image of M4 deletion (blue = DAPI, yellow = M1, red = M4). (**E**) Closeups showing M4 deletion from dorsal striatum while still intact in the nucleus accumbens, with accompanying expression percentages (n = 15). (**F,G**) Time course and averaged data showing a significant reduction in distance traveled in VU’154 group (unpaired t-test, p < 0.05). (**H**) Time course of change in dLight fluorescence in mCherry controls. (**I**) Averaged data showing no significant difference between groups (unpaired t-test, p > 0.05). (**J,K**) Time course and averaged locomotion data showing no significant difference between groups in DS-M4-KO mice (unpaired t-test, p > 0.05). (**L,M**) Time course and averaged AUC data of dLight in DS-M4-KO mice (unpaired t-test, p > 0.05).

Mice that were injected with AAV encoding mCherry demonstrated similar VU’154 effects on both AHL and amphetamine-induced DA release as was observed previously in wildtype (WT) mice (**Fig. 4F-I**). These include a significant attenuation of AHL (**Fig. 4F,G**; n= 5-7, * significantly different via unpaired t-test, p = 0.007) with a small elevation in amphetamine-induced DA release that did not reach statistical significance (**Fig. 4H,I**). However, mice injected with AAV encoding Cre-recombinase completely ablated VU’154-mediated reductions in AHL (**Fig. 4J,K**), with little to no effect on amphetamine-induced DA levels compared to vehicle-treated animals (**Fig. 4L,M**). These findings indicate that M4 receptors expressed in the DS are required for VU’154-mediated efficacy in AHL, yet do not robustly modulate amphetamine-induced DA release.

### VU’154-mediated decrease in spontaneous DA release amplitudes persists in DS-M4 KO mice

When examining VU’154-mediated effects on striatal DA under basal conditions, DS-M4-KO mice still exhibited a significant reduction in striatal DA release event amplitudes (**Fig. 5B-D**, n = 13, paired t-test, p = 0.001), similar to that of mCherry controls (**Fig. 5E,F**, n = 10, paired t-test, p = 0.04). As previously observed in WT mice, VU’154 did not significantly change the magnitude of DETQ effects on baseline fluorescence (**Fig. S5A,B**) but did reduce DA event amplitudes observed in the presence of DETQ (**Fig. S5C-H**, n = 11, p = 0.01 via paired t-test). Collectively, these results suggest that M4 receptors in the DS are not required to observe VU’154-mediated reductions in striatal DA event amplitudes.

**Figure 5:**
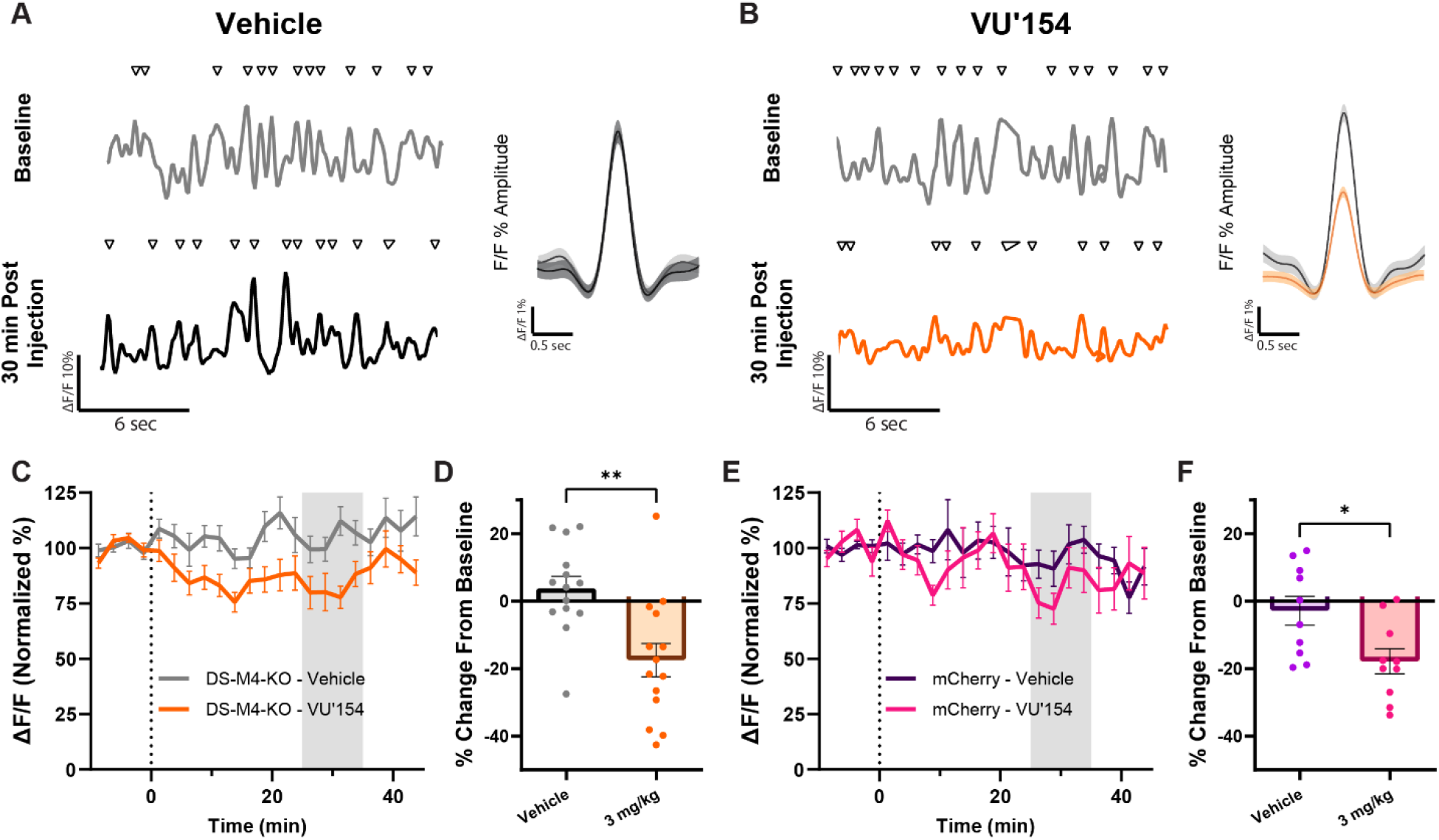
VU’154-mediated decrease in spontaneous DA release amplitudes persists in DS-M4-KO mice. (**A**) Representative snippets with detected spontaneous DA release events and mean composite trace with SEM 95% CI, at baseline and post injection, after vehicle and (**B**) 3 mg/kg of VU’154. (**C,D**) Time course and averaged data from mCherry controls depicting mean event amplitude (paired t-test, n = 10, p < 0.05). (**E,F**) Time course and averaged data from DS-M4-KO mice (n = 14, p < 0.005).

## Discussion

The antipsychotic potential of muscarinic compounds has advanced remarkably in a relatively short time, evolving from the serendipitous observation of xanomeline efficacy during Alzheimer’s disease clinical trials [42] to the ultimate approval of Cobenfy, the first non-dopamine receptor-targeting pharmacotherapy to be FDA-approved for schizophrenia [16]. Preclinical studies point to both M1 and M4 receptors as key mediators of these beneficial effects. The M4 receptor in particular is thought to play a key role in mediating antipsychotic-like effects, as supported by studies showing that highly subtype-selective M4 receptor PAMs can mediate antipsychotic-like efficacy in animal models that are correlated with clinical efficacy in treating positive symptoms [19, 26, 43–46]. However, the circuit-level mechanisms underlying M4 PAM-mediated antipsychotic efficacy *in vivo* remain incompletely understood. Consistent with previous reports [19, 39, 43], we confirm that the M4-selective PAM VU’154 robustly attenuates AHL. However, we found that VU’154-mediated effects on AHL are dissociated from concurrent changes in amphetamine-induced DA levels. We further demonstrate that VU’154 efficacy in AHL is absent in mice with a dorsal striatum-specific M4 deletion (DS-M4-KO; **Fig. 4J,K**), indicating that M4 receptors in the DS are required for antipsychotic-like efficacy.

### Dissociation of M4 PAM efficacy from amphetamine-induced DA release

The dissociation between VU’154 antipsychotic-like efficacy and amphetamine-induced DA levels was contrary to our initial expectations and differed from previous microdialysis studies done in the presence of mAChR-specific compounds and psychostimulants [19, 47, 48]. These studies differed from ours in technique (microdialysis vs. fiber photometry), species (mouse vs. rat), brain region (NAc vs. DS), stimulant administered (amphetamine vs. cocaine), and muscarinic drug administered (Oxotremorine vs. VU’100 vs. VU’154). Methodological differences cannot be ruled out, but we believe that they are unlikely to fully explain the discrepancy, as dLight-based fiber photometry has been reported to provide similar results to both fast-scan cyclic voltammetry [49] and microdialysis approaches for assessing amphetamine-induced changes in DA levels [34]. The majority of these previous studies were performed in the NAc, whereas we focused on the DS. This raises the possibility that mAChRs may differentially regulate DA release across different striatal subregions, and that M4 PAMs may have region-dependent effects on DA release in the NAc that are critical for M4 PAM efficacy in modulating substance use disorders involving alcohol [29] and cocaine [48]. Because DA signaling is most disrupted in schizophrenia patients in regions that are homologous to the DS of mice [8, 50], we created our DS-M4-KO to assess the importance of M4 receptors in the DS in mediating efficacy in AHL.

### Postsynaptic M4s on D1-SPNs of the dorsal striatum as a likely driver of antipsychotic-like efficacy

Our finding that VU’154 efficacy is absent in DS-M4-KO mice, together with previous results in mice lacking M4 in D1-expressing neurons [44], adds to evidence suggesting that postsynaptic M4 receptors expressed on D1-SPNs are a major driver of antipsychotic-like efficacy. M4 PAMs reverse not only stimulant-induced hyperlocomotion, but also D1-agonist-induced hyperlocomotion [44], which would be expected to occur independently of DA release. In addition, both xanomeline and VU’154 reduce Ca^2+^ transient rates in D1-SPNs, but not D2-SPNs, in the presence of amphetamine [46], while *ex vivo* slice studies indicate that M4 receptor activation attenuates D1 receptor-induced cAMP with no such effects on D2-SPNs [40, 51]. Given the lack of M4 on D2-SPNs [52], the latter studies further implicate a postsynaptic mechanism in lieu of alterations in extracellular DA. These findings, combined with imaging studies showing that M4 PAMs can reduce amphetamine-induced changes in brain activity [19], could suggest that M4 PAMs are able to suppress amphetamine effects on striatal function via postsynaptic effects that are independent of DA release mechanisms. However, future studies using cAMP imaging approaches in D1-SPNs *in vivo* will be needed to further test the hypothesis that M4 PAMs mediate antipsychotic-like effects through postsynaptic cAMP signaling.

### Effect of M4 activation on basal DA release

While M4 PAM administration had little to no effect on amphetamine-induced DA release, VU’154-mediated reductions in DA event amplitudes were observed under both basal and DETQ-enhanced conditions. DETQ is a PAM selective for the human ortholog of the DA D1 receptor (of which dLight is a modified form), with little to no activity at the mouse receptor. Thus, DETQ acts as a “potentiator” of dLight signals with minimal impact on brain circuit activity. In the absence of DETQ, chemogenetic inhibition of DA terminals was observed to have little to no effect on basal dLight1.3 signaling. However, in the presence of DETQ, the same chemogenetic intervention led to a significant inhibition of dLight signal [36], highlighting the utility of this tool compound in assessing the ability of interventions to reduce basal striatal DA levels. The inability of VU’154 to modulate baseline signals in the presence of DETQ suggests that there was little to no effect on tonic DA levels in the striatum. The persistence of VU’154-mediated reductions in DA event amplitudes in DS-M4-KO mice (**Fig. 5**) suggests that these effects are not mediated by M4 receptors in the striatum. These results implicate involvement of another population of M4 receptors, potentially located presynaptically on pedunculopontine and laterodorsal tegmentum (PPTg & LDTg) terminals to the substantia nigra pars compacta [53], on corticostriatal glutamatergic terminals [54], or potentially by M4 expressed on D1-SPN terminals in the substantia nigra pars reticulata [44]. As indicated by the persistence of M4 PAM-mediated reductions in DA transients in animals where antipsychotic-like efficacy was absent, this mechanism is likely less relevant to antipsychotic-like efficacy.

Nonetheless, future studies looking at region-specific deletion of these M4 receptor populations, or local infusions of M4 PAMs to these regions would be needed to further elucidate the mechanisms whereby M4 PAMs modulate DA event amplitudes *in vivo*. It is unclear why previous studies using brain slice preparations have demonstrated that mAChR activation can inhibit striatal DA signaling [26, 31, 55], while our results suggest that *in vivo* M4 modulation of DA release amplitudes is mediated by extra-striatal M4 receptors. One possible explanation is the need to use agonists when assessing PAM function in brain slice preparations, as tonic ACh levels in brain slices are not basally high enough to see PAM effects without experimental intervention to boost ACh levels [56]. Consistent with this idea, the efficacy of the mAChR agonist ML-007 in reducing AHL has been demonstrated to be equally dependent on both M1 and M4 receptors [21]. Accordingly, it is possible that M1 and M4 activation when combined may modulate DA release in ways that are distinct from when M4 alone is potentiated. Another potential contributor to these differences is that *ex vivo* studies assess either electrically-or optically-evoked DA release events, while our current study assessed spontaneously occurring DA release events. Follow-up studies assessing the ability of structurally distinct M4 PAMs and mAChR agonists (xanomeline) to modulate both spontaneous and evoked DA signaling *in vivo* are needed to further address these possibilities.

### Limitations

Several limitations should be noted. First, our DA release studies were performed in the DS (**Fig. 1C**, **Fig. 4C**), and our M4 deletion was broad across the DS (**Fig. 4B**) and they therefore do not differentiate between possible contributions of M4 in the lateral DS compared to the medial DS. Ultimately, future studies will be needed to assess the possibility that M4 expressed in the medial DS, lateral DS, and NAc may have differential effects on DA signaling or in mediating efficacy in attenuating AHL. Second, fiber photometry experiments measure bulk changes in DA signal, which may obfuscate small but physiologically relevant changes in local release dynamics that could play a role in mediating antipsychotic-like efficacy. To this point, the striatum exhibits both spatially restricted point-to-point transmission of DA, and bulk DA transmission, with emerging evidence demonstrating that these forms of neurotransmission can be differentially regulated and contribute to distinct behavioral outcomes [57]. Future studies using techniques to image DA dynamics with improved spatial resolution will be needed to address the ability of M4 PAMs to modulate spatially restricted DA transmission. Lastly, there is a need to assess M4 effects on DA release in other preclinical models with predictive validity, including assessment of DA during somatosensory gating [58], NMDA receptor antagonist-induced models [59], hallucination-like perception models [60], or models that selectively enhance DA in the DS [61], to gain a broader understanding of the relationship between M4 PAM antipsychotic-like efficacy and DA signaling.

## Conclusions

Taken together, these studies deepen our understanding of the mechanisms whereby muscarinic drugs produce antipsychotic-like efficacy. They highlight a key role for M4 receptors in the dorsal striatum and indicate that these effects are more complex than a broad normalization of amphetamine-induced DA release.

## Acknowledgments

The authors would like to thank Dr. Jürgen Wess for generously providing initial M4^fl/fl^ mice for our colony and Dr. Craig Lindsley for providing VU’154. Some of the figures were created with Biorender.com. We would like to thank Vianne Phan for tissue processing, and the Floyd School of Medicine Department of Laboratory Animal Resources for their work in maintaining the mouse colonies.

## Author Contributions

**DGJ**: Conceptualization, Methodology, Investigation, Data Analysis, Manuscript Writing and Editing. **CMJ**: Methodology, Data Analysis, Manuscript Editing. **YC**: Investigation and Manuscript Editing. **MNM**: Investigation and Manuscript Editing. **RDC**: Investigation and Manuscript Editing. **DJF**: Conceptualization, Methodology, Resources, Supervision, Manuscript Writing and Editing.

## Supplemental Methods

### Stereotaxic Surgery

Mice were anesthetized with an isoflurane-oxygen mixture (1.5-2.5% isoflurane) and mounted in stereotaxic apparatus (Stoelting). Viral vectors were injected using a Hamilton syringe. To secure cannulas, two screws, driven into pre-drilled holes, and a base layer of dental cement were applied to the skull for adhesion (Metabond powder, 1861068, Metabond base, 1864477, Amalgambond PLUS Catalyst, 1861119, Henry Schein), and the cannula was secured in place with Integrity dental crown material (Henry Schein 2226781). All mice received post-operative subcutaneous injections of carprofen at 50 mg/kg, with additional doses at 24 and 48 hours post-surgery. Mice were allowed 3+ weeks for healing and expression of virus before habituation and experimentation. At the end of experiments animals were sacrificed and brains were removed for immunohistochemistry analysis to confirm viral expression, cannula placement, and in some cases M4 muscarinic receptor deletion as outlined below. Any mice with insufficient viral transduction or cannula placement outside of the DS were excluded from analysis.

### Habituation

Prior to all experiments, mice underwent 3 consecutive days of habituation. Each day, mice were handled and weighed, connected to the fiber-photometry patch cable, administered a saline injection, and placed in the open-field arena for 10 min.

### Fiber Photometry

The setup utilized low-autofluorescence FC/PC and FC/PC-to-ferrule patch cables (Thorlabs) along with mirrorboxes and pig-tailed rotary joints (Doric).

### Assessment of Basal Striatal Dopamine Release and Locomotion

Dose-response experiments utilized a 10-min baseline prior to treatment injection, followed by 45 min of recording post-injection. Dosages given were 0, 0.3, 1, and 3 mg/kg. Tonic DA with DETQ experiments utilized a 30-min baseline for maximal photobleach stabilization for optimum normalization when calculating Z-score.

### Locomotion analysis

The DeepLabCut model was trained at 150,000 iterations. Pose-estimation data were filtered. Distance traveled was calculated from the centroid position, then analyzed using DeepLabCut-Analysis-Jupyter-Scripts, publicly available code developed by Farhan Augustine (10.5281/zenodo.10866995). Time in the center was defined as the time during which both the headbase and centroid were located within the central arena zone, defined as a circle centered in the arena with a radius equal to one-half of the arena radius.

### Tissue Processing

#### Immunohistochemistry

After perfusion with 4% PFA, brains were removed, post-fixed overnight in 4% PFA, and cryoprotected by sequential overnight incubation in 15% and 30% sucrose. Brains were sectioned on a freezing microtome. Sections were permeabilized in permeabilization buffer (0.5% Triton X100 in PBS) for 30 min prior to being incubated in blocking buffer solution (3% BSA, 3% normal goat serum, 0.3% Triton X100 in PBS) for 1 h. Sections were incubated overnight at 4°C with a chicken anti-GFP primary antibody (1:4000; ab13970, Abcam) in blocking buffer. The next day, sections were rinsed 3 times for 5 min in PBS and incubated for 1 h at room temperature in secondary antibody solution containing goat anti-chicken Alexa Fluor 594 secondary antibody (1:500; ab150176, Abcam) with Hoechst dye (2 μg/mL, Fisher). Afterward, sections were rinsed 3 additional times in PBS for 5 min, mounted with aqueous mounting medium, and dried overnight at 4° C. Images were acquired using an Invitrogen EVOS FL Auto Cell Imaging System.

### RNAscope

Mice were anesthetized in 2-3% isoflurane and euthanized by decapitation. Brains were rapidly removed, flash-frozen in 2-methylbutane, and stored at −80°C. Brains were subsequently sectioned coronally (20 µm) on a cryostat, collected on SuperFrost Plus slides (Fisher), and stored in a slide box that was pretreated with RNAse Zap (Fisher). Before the assay, slices were fixed for 1 h at 4°C in pre-chilled 4% PFA prepared in RNase-free PBS and then rinsed twice in RNase-free PBS. Sections were then dehydrated sequentially in 50%, 70%, 100%, and 100% ethanol. Probes mixtures were prepared according to the number of targets to be labeled using a 1x diluted probe mixture for Channel 1 as diluent as follows: M1 was labeled on Channel 1 (Mm-Chrm1-O1, 98%, ACD 483441), M4 on Channel 2 (Mm-Chrm4-C2, 2%, ACD 410581-C2), and ChAT on Channel 3 (, Mm-Chat-C3) were conjugated to, 2%, ACD 408731-C3), Fluorophores used are Channel 1: TSA Vivid 570 (yellow, 1:4000), Channel 2: TSA Vivid 650 (red, 1:5000), and Channel 3: TSA Vivid 520 (green, 1:3000). The assay was performed according to the manufacturer’s instructions. Following multiplex assay, sections were incubated in DAPI (RNAscope Multiplex V2 DAPI) for 30 s, and then mounted with ProLong Gold Antifade Mountant (Fisher) and cover-slipped. Sections are allowed to dry for at least 30 minutes and stored in the dark at 4°. Slides were imaged on the Leica Stellaris 5 LIAchroic Confocal System within 1 week of the assay.

### RNAscope image analysis

To measure ratio of M1-and M4-positive cells to total cells, M4, M1, and DAPI channels were converted to Luminance, then to RGB respectively in Fiji (Image J), and then analyzed in QuPath. QuPath’s Cell Detection was run on a non-deleted area (nucleus accumbens) with default values except for the following changes: Background Radius: 10 μm; Median Filter Radius: 2 µm; Sigma: 2 µm; Threshold: 1.

Custom code was used to optimize M1 and M4 puncta detection thresholds using the non-deleted image (https://doi.org/10.5281/zenodo.23301431), and these thresholds were utilized for puncta detection in the deleted area (dorsal striatum). Afterward the area of deletion was determined utilizing Fiji’s BigWarp plugin to fit fluorescent images to the corresponding atlas image and outlining the deletion.

**Supplemental Figure 1, related to Figure 1.**
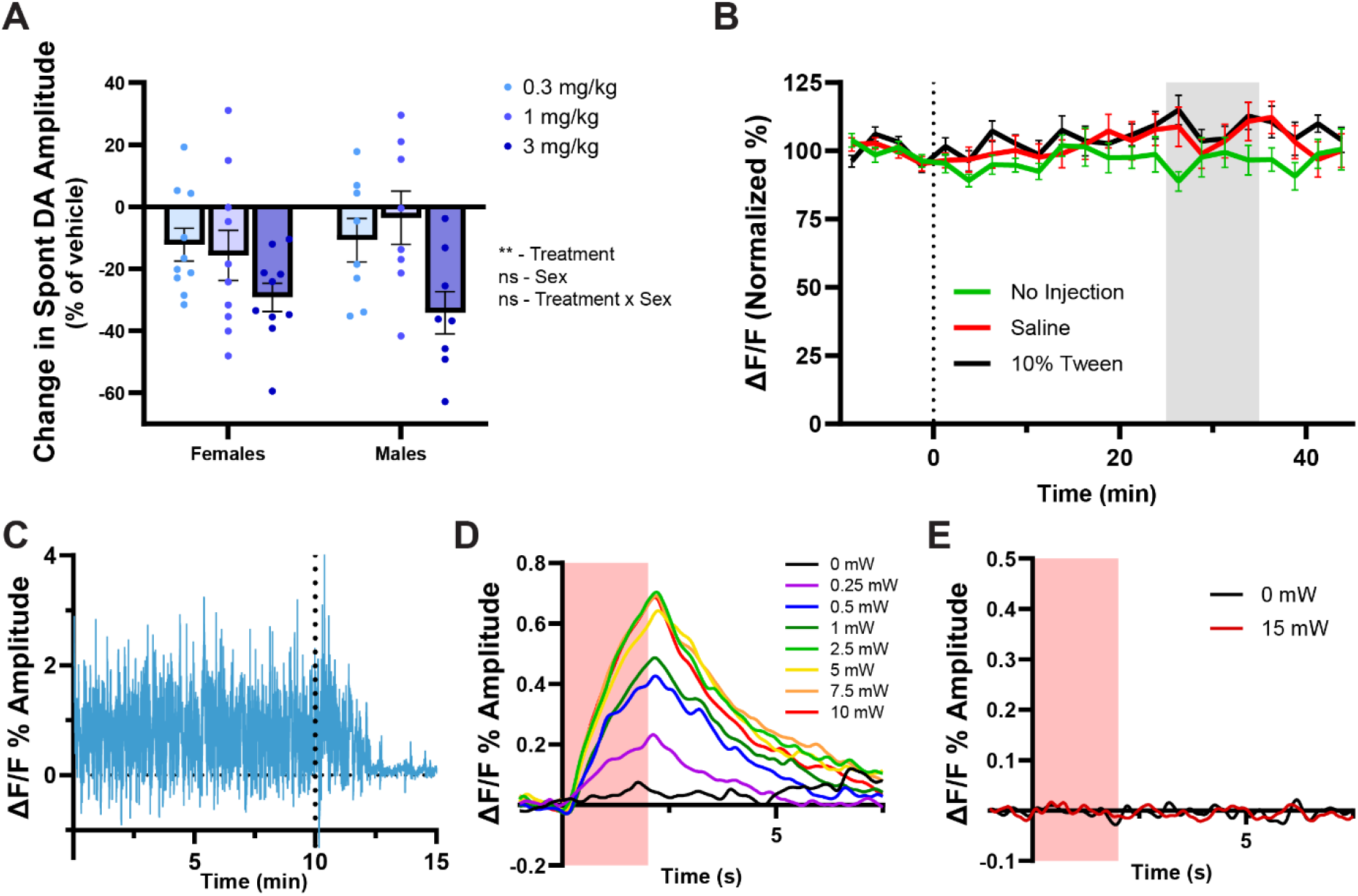
(**A**) Averaged data depicting change in amplitude comparing sexes with only a main effect of treatment (two-way ANOVA, treatment: p < 0.005). (**B**) Time course plotting event amplitudes vehicle (n = 17) to saline (n = 16) and no injection (n = 13) depicting apparent increased amplitude following injections. (**C**) Representative trial depicting s.c. administration of 10 mg/kg D1-like antagonist SCH-23390 (dotted line), confirming signal as dLight. (**D**) Input-output curve in DAT-Cre^+/−^mice expressing dLight along with Chrimson (red portion = 20 Hz pulse duration). (**E**) Time course of optogenetic pulse applied in mice expressing dLight without Chrimson.

**Supplemental Figure 2, related to Figure 2.**
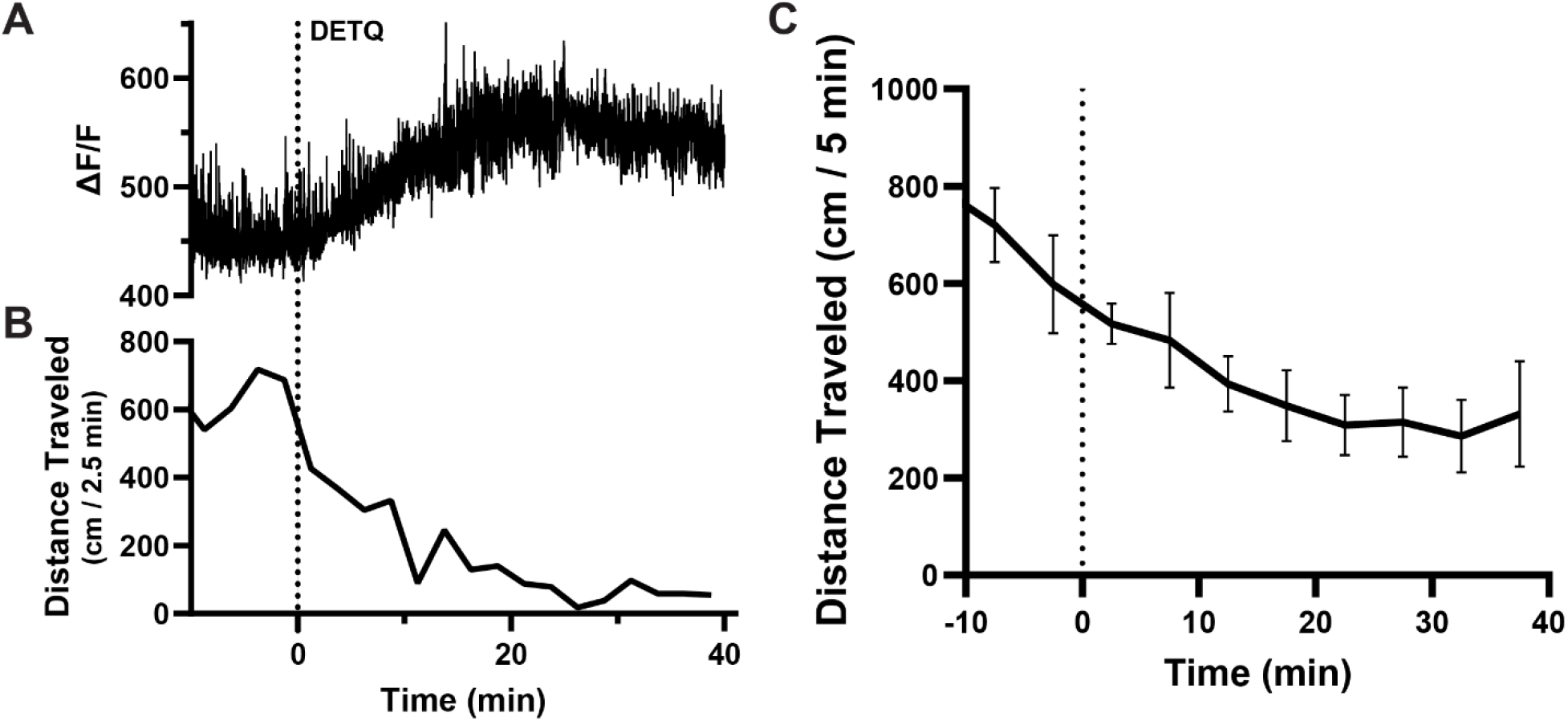
(**A**) Representative trial depicting change in dLight fluorescence before and following DETQ administration (10 mg/kg; i.p.). (**B**) Representative trial depicting locomotion change over time, before and after DETQ. (**C**) Time course of distance traveled prior to and after DETQ administration with vehicle (10% Tween 80; i.p.).

**Supplemental Figure 3, related to Figure 3.**
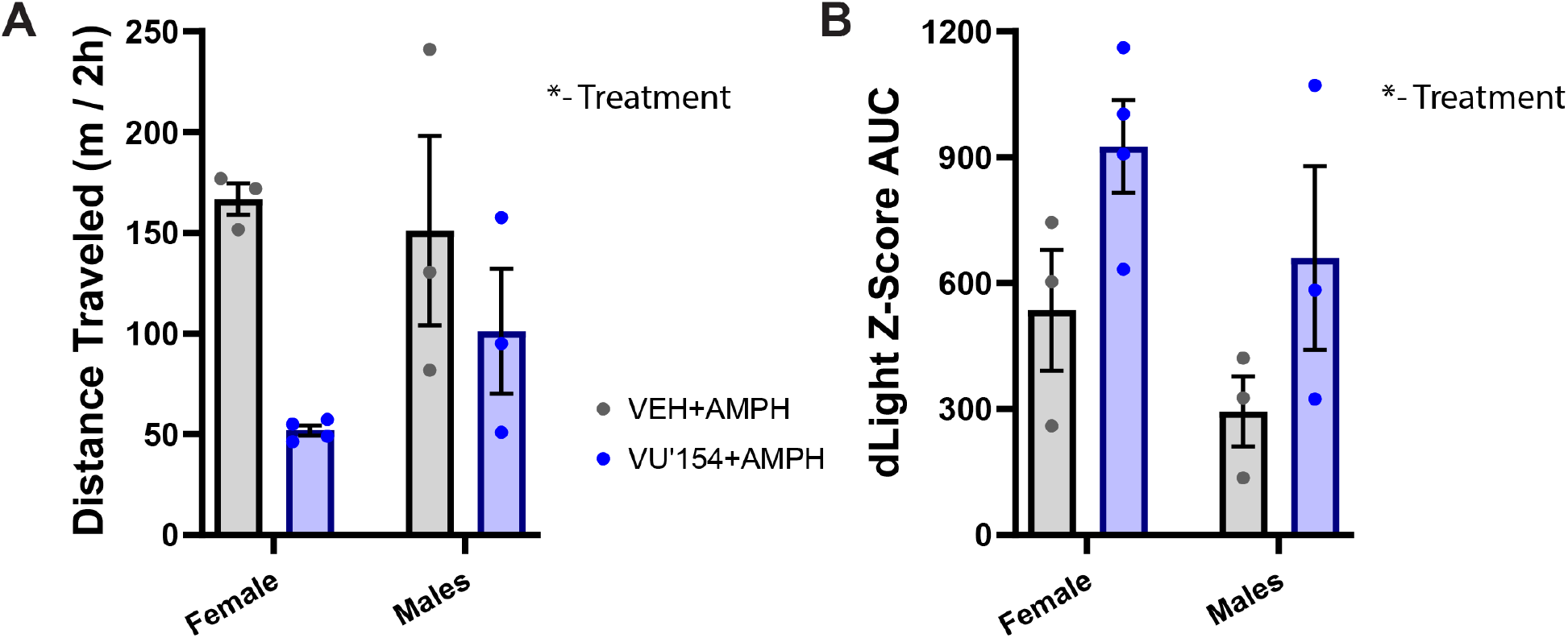
(**A**) Averaged data depicting distance traveled post-amphetamine split by sex, which exhibited a main effect of treatment (p = 0.012), but no effect of sex (p = 0.535). (**B**) Averaged data depicting dLight Z-score AUC post-amphetamine, which showed a main effect of treatment (p = 0.028), but no effect of sex (p = 0.113)

**Supplemental Figure 4, related to Figure 4.**
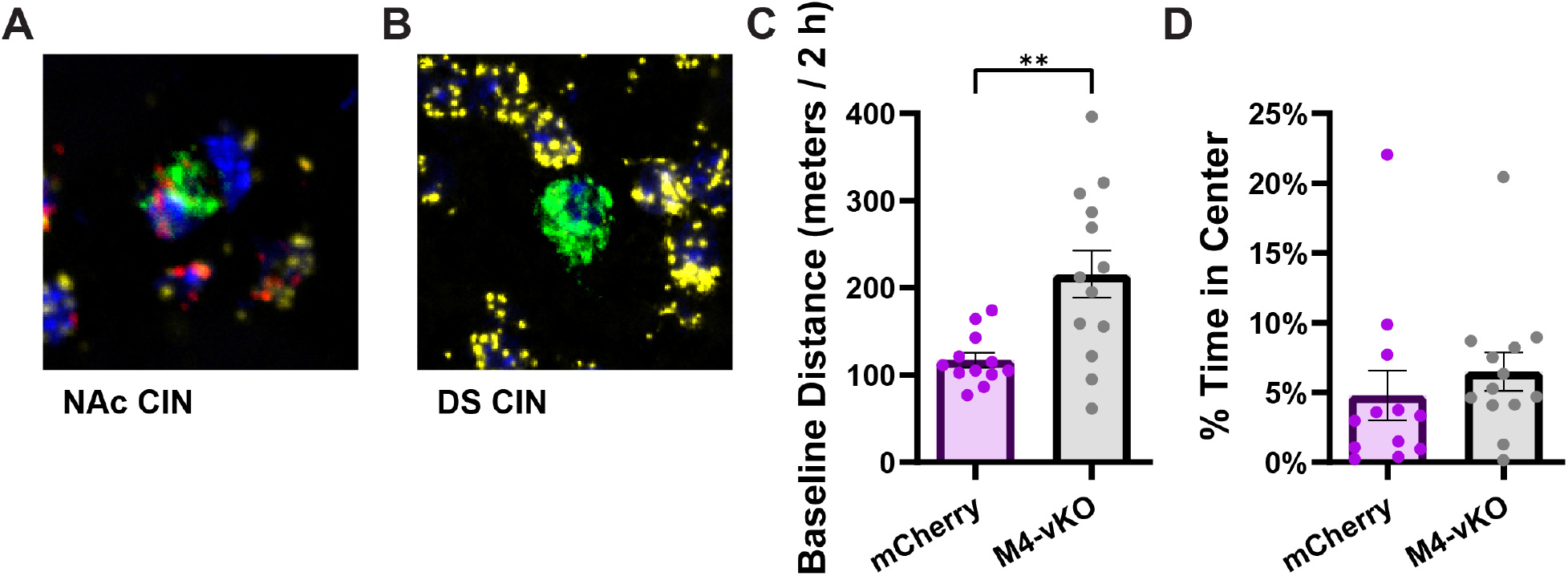
(**A**) Representative image of cholinergic interneuron in the NAc expressing M4 (red). (**B**) CIN in the DS with no M4 present. (**C**) Averaged data (mean ± SEM) comparing distance traveled prior to drug administration (unpaired t-test, p < 0.005). (**D**) Averaged data comparing time spent in center prior to drug administration (unpaired t-test, p > 0.05).

**Supplemental Figure 5.**
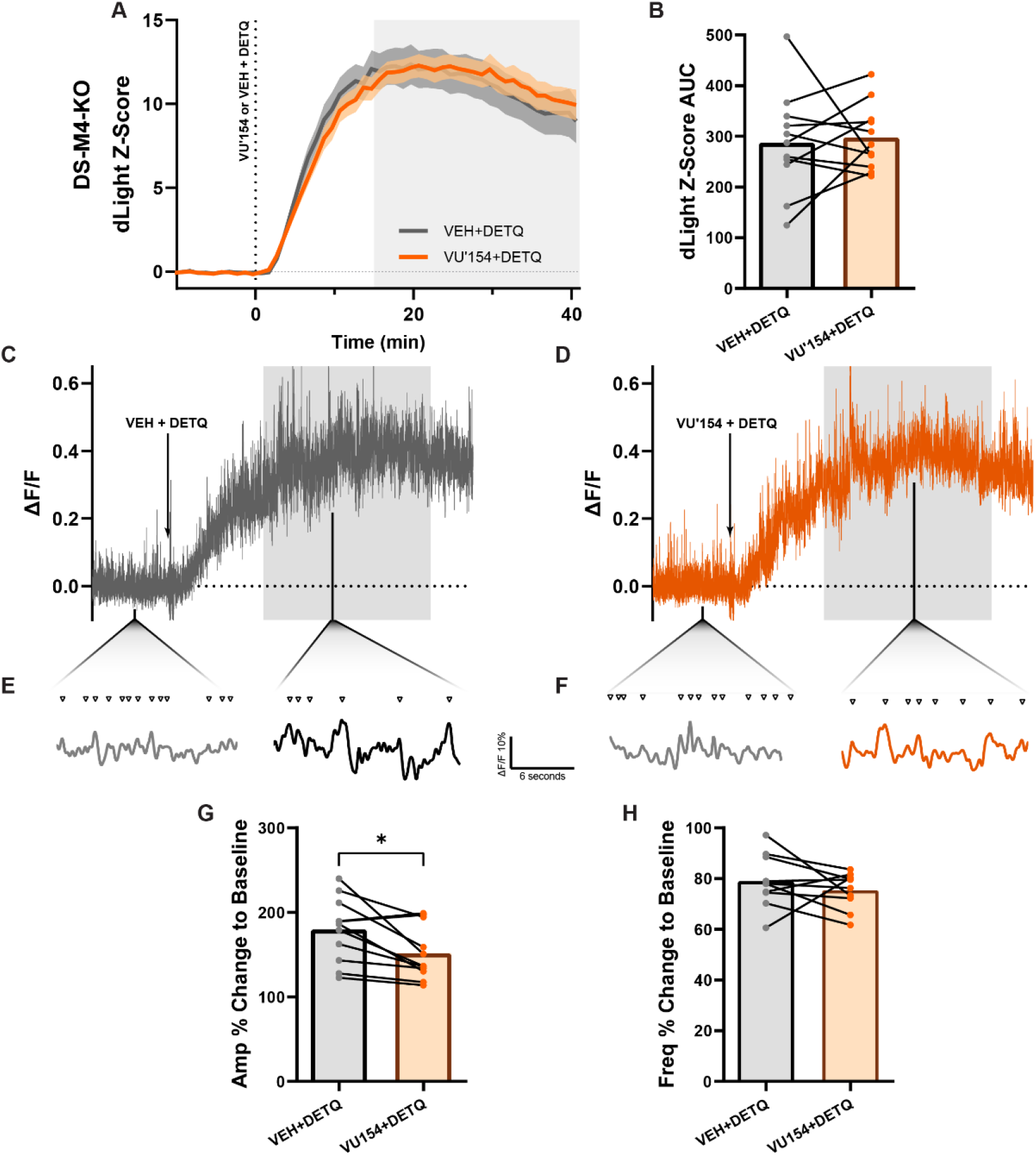
(**A**) Time course of change in measured dLight fluorescence intensity (Z-score relative 10 min pre-injection. (**B**) Averaged data depicting dLight Z-score AUC (paired t-test, p > 0.05). (**C**) Representative trial trace following coadministration of DETQ (10 mg/kg i.p.) with vehicle and (**D**) 3 mg/kg of VU’154. (**E**) Snippets of trace labeled with detected events and base line and peak period after DETQ with vehicle and (**F**) 3 mg/kg of VU’154. (**G**) Averaged data depicting mean event amplitudes at peak period relative to baseline (within subjects) (n = 10, paired t-test, p < 0.05). (**H**) Averaged data depicting mean event frequency at peak period (paired t-test, p > 0.05).

